# Robust maintenance of both stimulus location and amplitude in a working memory model based on dendritic bistability

**DOI:** 10.1101/2025.07.07.663443

**Authors:** Jiacheng Xu, Daniel L Cox, Steven J Luck, Mark S Goldman

**Affiliations:** Department of Physics and Astronomy, University of California Davis, Davis CA 95616 USA; Center for Mind & Brain and Department of Psychology, University of California Davis, Davis CA 95616 USA; Center for Neuroscience, and Department of Neurobiology, Physiology, and Behavior, University of California, Davis, Davis, CA 95616, USA; Department of Ophthalmology and Vision Science, University of California Davis, Davis CA 95616 USA

## Abstract

Working memory is a core feature of cognition that enables items to be maintained and manipulated over short durations of time. Stored information can be binary, such as the presence or absence of an object, or graded, such as the graded intensity or location of a feature. Current computational models of working memory cannot robustly maintain both the graded intensity and spatial location of a stored item. Here, we show how this limitation can be overcome if neurons contain multiple bistable dendritic compartments. First, we illustrate the core mechanism for the storage of graded amplitude information in a simple spiking “autapse” circuit consisting of a single neuron connected to itself. Second, we reduce this model to a rate-based model that permits analytic understanding. Third, we implement this mechanism within a spatially extended architecture in which the spatial location of an item is encoded by the set of active neurons. In contrast to classic spatial working memory models, which only encode the binary presence of an item at a given location, the multi-dendrite-neuron model robustly encodes both the amplitude and location of an item in working memory in a noise-resistant manner and without requiring fine tuning of parameters. We show analytically that the key mechanism permitting the storage of amplitude information is equivalent to that of the simpler autapse circuit. This work provides a solution to the problem of encoding graded information in spatial working memory and demonstrates how dendritic computation can increase the representational capacity and robustness of working memory.

**Significance Statement:** Animals can readily remember both the location of an item and analog features of the item such as its amplitude or intensity. Remarkably, current computational models of working memory require extreme fine tuning of model parameters to perform this task. Here, we show how this limitation can be overcome if neurons have active dendritic processes that enable local dendritic compartments of the neuron to robustly maintain digital “up” (“plateau potential”) or “down” states. By activating a variable number of plateau potentials at each location, the models can jointly maintain both the location and amplitude of a stimulus in working memory in a noise-resistant manner. This work demonstrates how local dendritic processes can enhance the computational capabilities of working memory networks.

## Introduction

Working memory refers to a mechanism that provides a temporary storage buffer for cognitive tasks such as search, reasoning, and decision-making. Working memory often must precisely store analog quantities such as the location, amplitude, or color of an object. The mechanisms by which such analog quantities are stored robustly in working memory has been puzzling, as the continuous attractor mechanisms traditionally used to describe the dynamics of analog working memory require fine tuning to avoid instability and are subject to drift in the presence of noise [1, 2, 3, 4, 5].

Classic models of graded working memory are based on line attractors. In a line attractor, stored memories are encoded into a continuous set of stably maintainable neural activity states, where each state represents the value of a distinct analog feature. Such models typically fall into two classes. In one class, which we term “location-coding” [2, 6], the encoded quantity is represented by a fixed amplitude pattern of neural activity that can shift between different neural locations, with the location of the pattern representing the analog stimulus [7]. This class of models has commonly been used to describe the encoding of variables such as the spatial location of an object [8], the orientation of an animal’s head in space [9], or the location (hue) of a color on a color wheel [10]. The canonical model of this class is the “ring” model in which neurons are arranged in a one-dimensional ring, with each neuron preferentially encoding a particular location in a circular stimulus space (Fig. 1A). This model has found direct experimental support in the microscopic connectivity of the fly head direction system [11] and has been widely used to explain many cognitive tasks [12].

**Figure 1:**
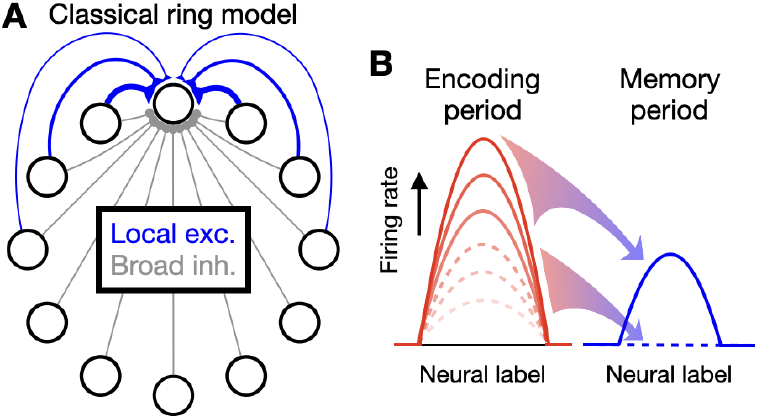
Schematic overview of inability of classic ring models to memorize graded amplitudes. **A**, Classical ring model. Neurons are arranged in a one-dimensional ring with local excitation between nearby neurons (blue) and broad inhibition across all neurons (gray). **B**, Inability to memorize graded amplitudes. In classical ring models, only two stable memory states exist: a fixed amplitude pattern of activity (solid blue line) that occurs when the encoding period neural activities, caused by different stimulus amplitudes, have high firing rates (solid red lines), or an untuned baseline level of activity (dashed blue line) that occurs when the encoding period activities have low firing rates (dashed red lines). Graded values of stimulus amplitudes are not memorized.

The second class of models, which we term “amplitude-encoding” [2, 3, 13], encodes memorized quantities in the amplitude of a largely fixed spatial pattern of neural activity. This class of models has been used to represent the memory of amplitude-or intensity-like variables such as luminance [14] or vibration frequency [15]. Such models have direct anatomical support in the connectome of the oculomotor integrator of the eye movement system [16] and are the canonical neural implementation of accumulation-of-evidence based decision-making [17, 18].

In actual organisms, working memory can simultaneously encode both the location and amplitude of an item. Further, even when encoding only the location of an item, Bayesian theories posit that the amplitude of the spatial pattern of neural activity represents the precision with which the location is encoded [19, 20]. Despite this, few models exist of such joint amplitude-location encoding, and the few existing models exhibit a lack of robustness in maintaining analog amplitudes of neural firing. This lack of robustness reflects fundamental limitations of classic models of location and amplitude encoding. Classic location-coding models like the ring model exhibit only a single all-or-none “bump” of activity and thus do not represent amplitude information (Fig. 1B). Further, they require either a perfectly symmetric network connectivity [7] or a functionally similar connectivity [4, 21, 22] to maintain a graded representation of location. Classic amplitude-coding models require fine tuning of their recurrent synaptic connectivity to achieve stable encoding of analog stimulus amplitudes – recurrent feedback that is too weak or too strong leads to rapid decay or growth, respectively, of neural activity on the timescale of the intrinsic neuronal or synaptic time constants of the model. Previous models of joint amplitude-location encoding have inherited these shortcomings and either require fine tuning of parameters to store analog amplitudes [23, 24, 25, 26] or trade off sensitivity in representation against robustness [27]. Further, even when fine-tuned, such models are subject to rapid diffusion of representations in the presence of noisy neural activity.

A separate set of amplitude-coding memory models has succeeded in overcoming the fine tuning problem by incorporating bistable memory components. In these models, analog memory values are generated through summing up the contributions of many bistable neurons [13] or dendritic compartments [28, 29]. These models inherit the robustness of the bistable processes upon which they are built, while also being capable of storing a graded representation of amplitude limited only by the number of bistable components. However, in location-coding memory models, only cellular bistability has been employed [30, 31], which stabilizes the memory of a given location, but does not permit the joint memorization of stimulus amplitude. Biophysically, bistability can be generated by positive feedback among small clusters of neurons [13] or, at a cellular level, through positive-feedback interactions mediated by the ionic conductances that underlie somatic [32, 33, 34, 35, 36, 37, 38, 39, 40] or local dendritic [41, 42, 43] plateau potentials.

Here, we show how a network of neurons with bistable dendrites provides, to our knowledge, the first model that jointly and robustly encodes location and amplitude information. We first demonstrate the core amplitude-storage biophysical mechanism through construction of a spiking, multi-dendrite “autapse” model consisting of a single neuron connected to itself [1]. We then simplify this biophysical model to an analytically solvable firing rate model that provides rigorous insight into the mechanism of amplitude storage. We next construct a multineuron ring model and show how, through a mechanism that can be directly mapped onto that of the simple autapse model, the network can represent both the analog amplitude and analog location of a stimulus. The model exhibits robustness to noise in its encoding of both memory amplitude and location, thus demonstrating how bistable neuronal processes can enhance the performance of working memory.

## Materials and Methods

Here we provide the equations and numerical methods underlying the three models described in the main text. First, we present the equations and simulation methods for the spiking autapse circuit model. Second, we present the equations governing the dynamics of the rate-based autapse circuit model, including those that describe the multistable band that geometrically determines the model’s stability properties. Third, we present the dynamical equations for the multistable ring model. Finally, we present numerical simulation specifications for the models, including characterizing the ring model’s behavior in the presence of noise. Mathematical derivations of the exact or approximate behaviors of the rate-based models are provided in the Mathematical Appendix.

### Spiking autapse circuit model with multiple conductance-based, bistable dendrites

The spiking autapse circuit model consists of an integrate-and-fire soma with N_d_ = 10 conductance-based dendrites. The somatic voltage V_s_ is governed by

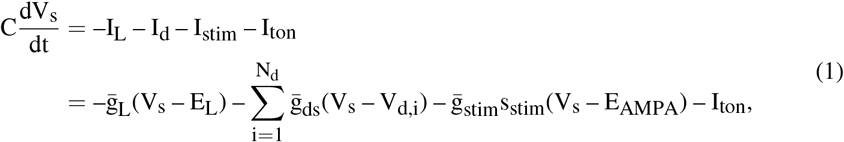

where I_L_ is the leak current, I_d_ is the summed current received from all dendrites, I_stim_ is the excitatory current representing the external stimulus, which arrives through AMPA receptors, and I_ton_ is the tonic background current. We chose parameter values similar to those of a previous model [43]. The membrane capacitance is C = 10 nF/mm^2^. The maximal conductances are 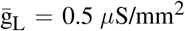 and 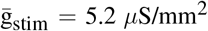. The conductance mediating the flow of currents from dendrite to soma is 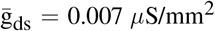. The equilibrium potentials are E_L_ = –80 mV and E_AMPA_ = 0 mV. The tonic current is I_ton_ = –17.2 nA/mm^2^. The synaptic activation variable s_stim_ obeys dynamics described below. The firing threshold for the soma is V_th_ = –50 mV. Each spike sets the somatic voltage to 30 mV for 3 ms, after which it resets the somatic voltage to -80 mV.

The voltage dynamics of the i^th^ dendritic compartment, V_d,i_, were adapted from [43], with modified maximum conductance values and the inward rectifier potassium (Kir) conductance simplified to not include a synaptically driven component:

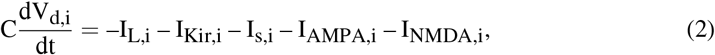

where the first two terms give the intrinsic dendritic leak current I_L,i_ and inward-rectifying potassium current I_Kir,i_, the third term I_s,i_ gives the current arriving from the soma, and the final two terms give the AMPA and NMDA components of the synaptic currents driven recurrently by somatic spiking. These currents are described by the following equations:

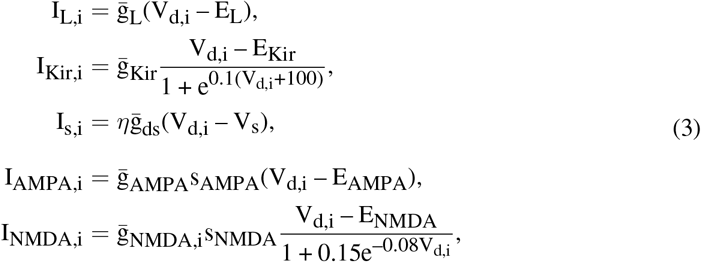

where *η* = 2 represents the ratio of the area of the soma to that of each dendrite. E_NMDA_ = 0 mV, E_Kir_ = –90 mV, 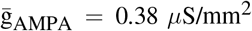 and 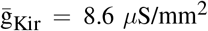. The maximum conductance for the NMDA receptors is 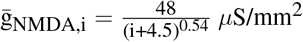.

The synaptic activation s_*α*_ obeys:

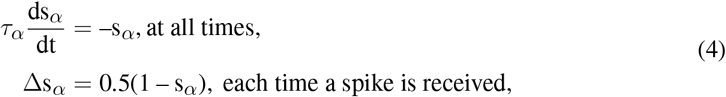

where *α* denotes AMPA (s_stim_ and s_AMPA_) or NMDA (s_NMDA_) receptors with a decay time constant *τ*_AMPA_ = 2 ms or *τ*_NMDA_ = 100 ms, respectively.

To derive the individual dendrite bistable input-output relations shown in Figure 2B, we removed all of the self-connections shown in Figure 2A. Next, a periodic train of spikes whose firing frequency steps up and down across a wide range of firing frequencies was delivered to each dendrite while its dendritic voltage was recorded. During this process, the dendritic voltage is also affected by somatic voltage changes through the electrical coupling between the somatic and dendritic compartments. To assure that the dendritic input-output relation was assessed with realistic somatic voltage levels, we confirmed that the somatic spike train driven by the dendritic inputs resembled that of the periodic inputs delivered to the dendrites. For the simulation, V_s_ and V_d,i_ were initialized to –70 mV, and the periodic train of input spikes delivered to each dendrite started at a rate of 0 Hz. The spike train lasted for a period of 1400 ms, and the averaged dendritic voltages over the last 1000 ms were recorded for all dendrites simultaneously. Next, the frequency of the periodic input train was increased by 1 Hz, and the recording repeated. Once the frequency reached 70 Hz, it was decreased by 1 Hz at each recording until returning to 0 Hz.

**Figure 2:**
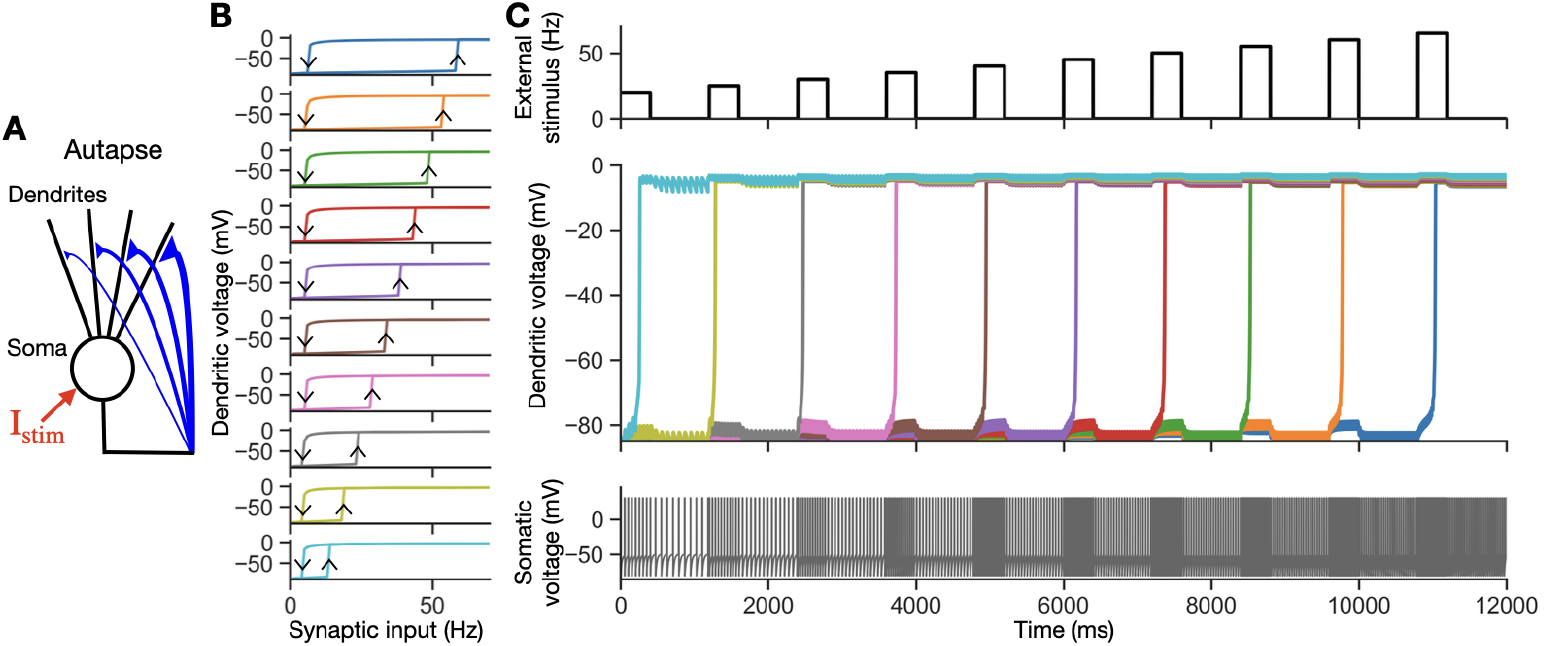
Maintenance of graded firing rates in the spiking autapse model. **A**, Model architecture. The model consists of an integrate-and-fire soma that projects to ten conductance-based dendritic compartments with bistability based on the voltage-dependent properties of the NMDA and Kir channels (see Materials and Methods). Each compartment receives synaptic input with a different weight (the maximal NMDA receptor conductance, 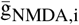), so that different dendritic compartments are recruited sequentially with increased amplitude of synaptic input. The external stimulus I_stim_ (red) goes into the soma. **B**, Hysteretic input-output relations of the ten dendrites. To characterize the bistable input-output relations of each dendrite, the synaptic inputs from the soma were removed and replaced by periodically firing, externally applied synaptic inputs that stepped across a range of different frequencies (x-axis). The voltages shown represent averages over 1000 ms of recording. Arrows in the top panel indicate the directions of the hysteretic response. Top to bottom panels show the 10 dendrites, ordered by increasing NMDA conductance 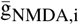 (or equivalently, decreasing up threshold). **C**, Memorization of different amplitudes. For the model in **A**, an external stimulus spike train to be encoded by the circuit is briefly applied, followed by a memory period with no stimulus. This is repeated 10 times for successively increasing external stimulus frequencies (top). Dendrites are activated during the encoding periods, starting from the one with the largest 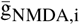, and self-sustain their activity during the memory period (middle). Somatic activity, which represents the memorized stimulus during the memory period, reflects the number of activated dendrites (bottom).

To demonstrate that the circuit maintains multiple levels of activity (Fig. 2C), we implemented the following procedure: V_s_ and V_d,i_ were initialized to –70 mV, and the circuit dynamics were allowed to stabilize for a period of 1200 ms before providing a sequence of external stimuli. Each external stimulus in the sequence consisted of a step on and off of periodic spikes for 400 ms, followed by an 800 ms memory period with no external stimulus. The frequency of the external stimulus sequence started at 20 Hz and increased by 5 Hz for each step until reaching 65 Hz. For the parameter values given above, this procedure activated one additional dendrite with each step increase in frequency, as shown in Figure 2C.

### Rate-based autapse circuit model

#### Simplified autapse dynamics

The dendrites in the rate-based autapse circuit model were modeled with a simple bistable input-output relation that approximated that observed in the biophysical model. Specifically, each dendrite has a bistable input-output relation as a function of the firing rate r of the form (Fig. 3A):

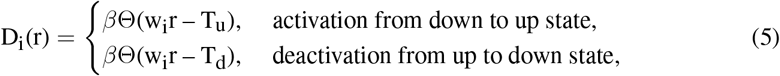

where *β* = 1 is the dendritic contribution to firing rate for an up-state dendrite, Θ(x) indicates the Heaviside (unit step) function, w_i_ is the weight of the connection onto the i^th^ dendrite, and T_u_ = 9 and T_d_ = 2 are constant up and down thresholds. Dendrites are ordered such that w_i_ is a monotonically decreasing function of the dendrite index i. A larger synaptic weight w_i_, which corresponds to a larger value of 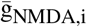 in the spiking model, effectively lowers the up and down thresholds. We can then rewrite D_i_(r) in terms of threshold firing rates for activating or deactivating a dendrite,

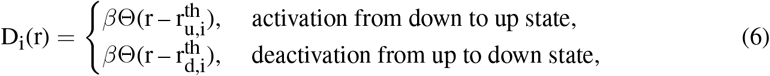

where the effective up and down thresholds are:

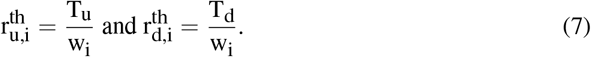

**Figure 3:**
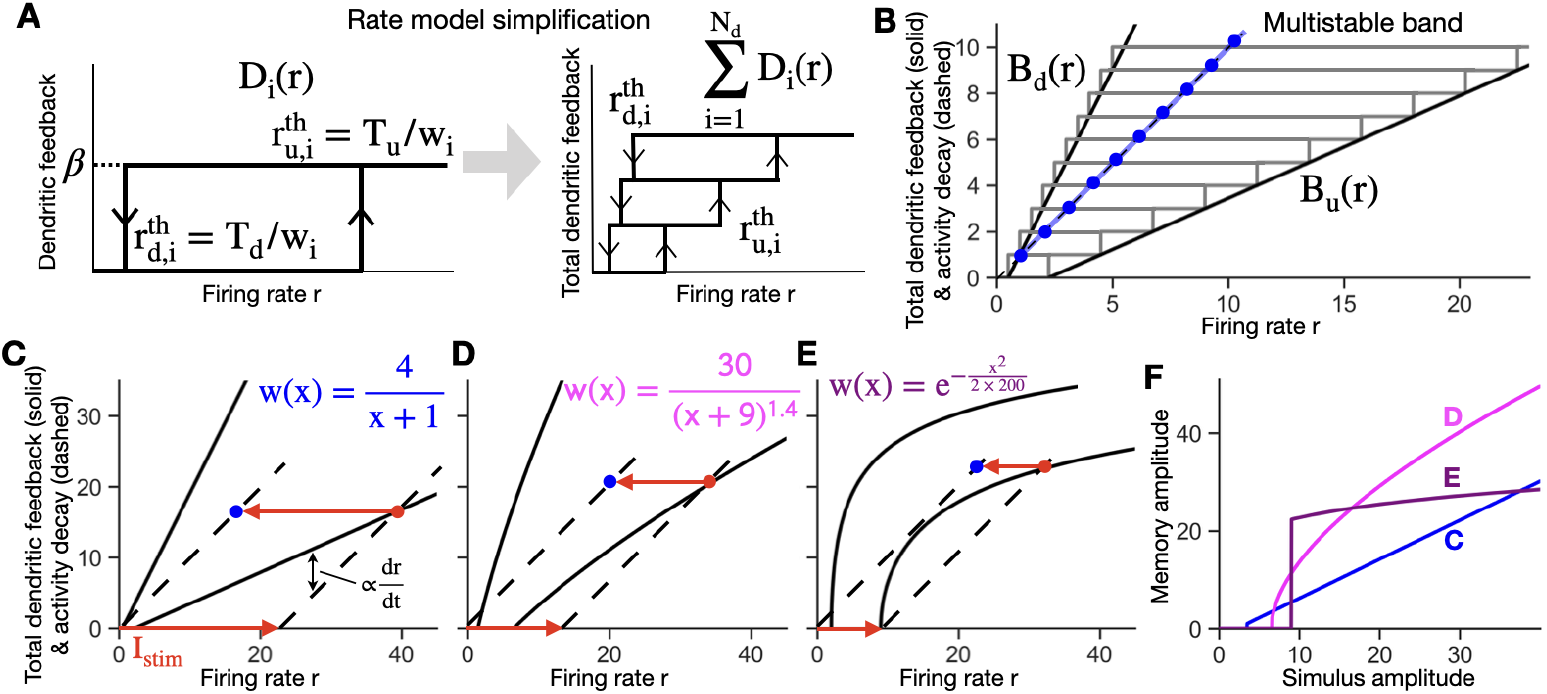
Rate-based autapse model for storage of graded amplitude stimuli. **A, B**, Rate based model captures the key behavior of the spiking model of Figure 2B. **A**, Left, bistable input-output relation D_i_(r) of a dendrite i. The input is the somatic firing rate, which feeds back to the dendrite through the self (autaptic) connection. D_i_(r) is hysteretic with jumps at up and down thresholds, whose values are inversely related to the synaptic weight w_i_. Right, summed dendritic output to the soma from all dendrites (3 shown). Each rectangle represents the contribution from one dendrite. Dendrites with smaller w_i_ values have smaller thresholds and are plotted at the bottom. **B**, Multistable band in the discrete or continuous case. Multiple dendrites form a band with right and left edges B_u_, B_d_ consisting of up and down thresholds, either in discrete steps (gray) or the continuum limit (black). The dashed line represents the decay of firing rate activity (magnitude of term –r in Eq. (15)). Intersections of the line and multistable band (the blue line in the continuous case or blue dots in the discrete case) represent points at which the total dendritic feedback offsets firing rate decay to lead to persistent firing (i.e., memory) states in the absence of an external stimulus. **C-E**, Different weight functions lead to different shapes of the multistable band and different responses to an external stimulus. **C**, Linear band derived from a power law decay weight function with power p = 1 (Mathematical Appendix). During the encoding period, an external stimulus I_stim_ shifts the dashed line (representing the combined effect of neuronal decay and the external stimulus) rightwards, with activity stabilizing at the red dot. The rate dr/dt of firing rate increases is proportional to the difference between the dashed line and rightward band edge B_u_. In the memory period following the offset of the external stimulus I_stim_ (red leftward arrow), memory activity stabilizes at the blue dot. **D**, Same as **C**, but for a power law weight decay with p = 1.4. **E**, Same as **C**, but with an example Gaussian weight function. For this case, when the external stimulus barely exceeds the smallest up threshold, the memory activity abruptly jumps to a high level, amplifying the external stimulus. **F**, The form of the weight function determines the shape of the relationship between the stimulus amplitude and the amplitude of stored memory activity. Traces show the mappings between stimulus and memory amplitude for each weight function used in **C-E**.The blue trace represents linear encoding of stimuli in memory (Mathematical Appendix).

For analytic simplicity, we assumed that the somatic firing rate dynamics are given by an exponential approach to a steady state given by the sum of the dendritic contributions and the input I_stim_ that represents the strength of the stimulus to be remembered by the circuit:

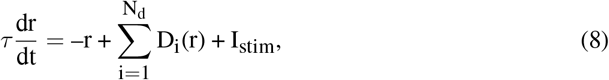

where N_d_ is the number of dendrites. *τ* is chosen to capture, in a simple manner, the slowest timescale of neural activity, which is set by the NMDA receptor kinetics in the spiking model. Explicit inclusion of additional synaptic or dendritic dynamics does not qualitatively change the primary results of this work.

#### Determination of the shape of the multistable band

The total dendritic feedback to the soma 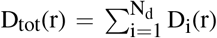 graphically defines a band of multistable memory storage. Here, we provide the methods by which the edges of this band and their continuum approximation (Fig. 3) are defined. Following Eq. (8), the right side of the band, which consists of the set of up thresholds of sequentially activated dendrites, is given by

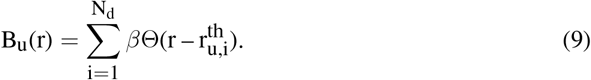

For a large number of dendrites, the weight function w_i_ and the edge of the band can be approximated by a continuous set of values, enabling the band to be described by a continuous functional form. Replacing the discrete index i by a continuum index x ∈ [0, N_d_] and assuming w_i_ changes gradually, we approximate the discrete set of w_i_ by a continuous monotonically decaying function w(x). Then we have:

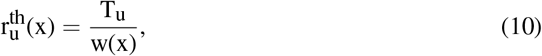

and B_u_ becomes:

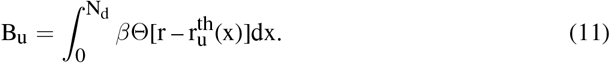

To derive an analytical expression for the right side of the band, we evaluate the above integral. The integrand is zero for 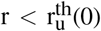 and a constant value of *β*N_d_ for all values of x such that 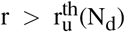. For intermediate firing rate values, using Eq. (10), every dendrite with an x value satisfying r > T_u_/w(x) contributes to the integrand, so that the total number of activated dendrites equals w^−1^(T_u_/r). Thus, the values of the right side of the multistable band, B_u_(r), correspond to *β* times the number of dendrites activated at a steady-state firing rate r:

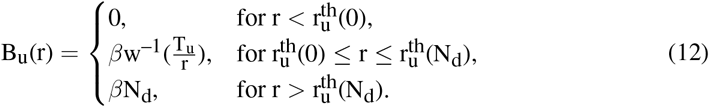

Similar reasoning applies for deriving the left side of the band, B_d_(r).

### Rate-based model with ring architecture

The ring architecture network underlying Figures 4-7 and illustrated in Figure 4A was constructed as follows. The network consists of N neurons, with each neuron having N_d_ = N dendrites. As in traditional ring models, except in Figure 7C, the recurrent excitatory weights w_ij_(d) were chosen to be symmetric, with a falloff in strength as a function of the distance d between neurons along the periodically arranged ring. The recurrent inhibitory weights were chosen to be global and of uniform weight w_inh_. Each excitatory connection projects to a separate bistable dendrite on the postsynaptic neuron, and each inhibitory connection as well as the external stimulus project to the soma of the postsynaptic neuron. The firing rate dynamics of each neuron is given by

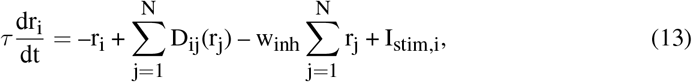

where D_ij_(r_j_) is the bistable dendritic input-output relation, and I_stim,i_ is the external stimulus input. The dendritic relation D_ij_(r_j_) for the contribution to the firing rate of neuron i from the dendrite receiving input from neuron j, is defined analogously to D_i_(r) in the rate-based autapse model:

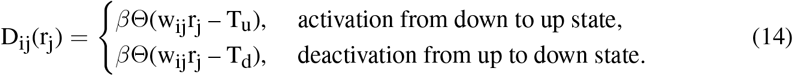

**Figure 4:**
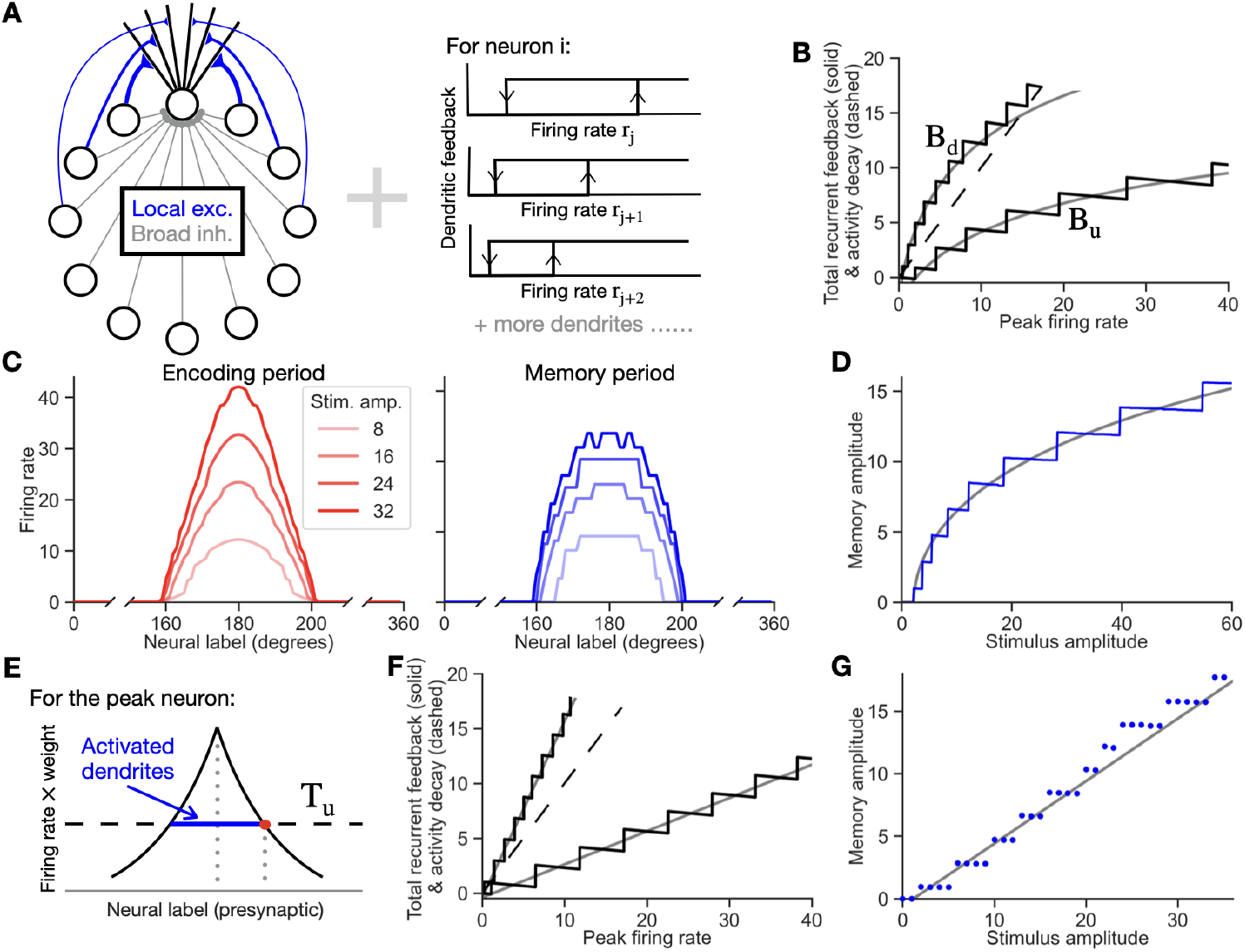
Simultaneous storage of stimulus location and amplitude in the ring model with dendritic bistability. **A**, Network architecture. Left, each neuron receives local excitatory input through connections onto separate dendrites and broad (uniform) inhibitory input through connections onto the soma. Right, each dendrite is bistable. Dendrites receiving inputs through larger synaptic weights (bottom) have lower thresholds for presynaptic neuron firing to flip the dendrite to its up state. **B**, Simulated multistable feed-back band for peak firing neuron (black) and approximate, analytically calculated multistable feedback band for the whole ring model network (gray). See Materials and Methods for details. **C**, Storage of stimulus amplitude at a given location. Left, steady-state pattern of activity during the encoding period for a stimulus with the shown amplitude (legend) and location centered at 180 degrees. Right, stable pattern of persistent activity during the memory period. Each stimulus amplitude is faithfully encoded during the memory period. **D**, Mapping between stimulus and memory amplitude. Blue, simulations. Gray, approximate, analytically calculated relationship (see Mathematical Appendix for derivation). **E**, Schematic illustrating how different presynaptic inputs contribute to a postsynaptic neuron’s level of activity during the encoding period (see Mathematical Appendix). The effective input a postsynaptic neuron receives at each dendrite is given by the product of the presynaptic neuron’s firing rate and synaptic weight onto the dendrite. If this product exceeds the up threshold value T_u_ (dashed line), the dendrite is activated. The total number of activated dendrites of a neuron equals the length of the blue segment. The example shown is for the peak neuron at the center of a local stimulus pattern. **F**, Multistable band for a network with weights chosen to produce a linear relation between stimulus and memory amplitude. Black, simulated multistable band for the peak neuron. Gray, approximate analytically calculated band for the whole network. **G**, Mapping between stimulus and memory amplitude for weights analytically calculated to approximate a linear relation, as in **F**.

#### Simulation details in the noise-free case

For all ring network model simulations, we simulated N = 360 neurons, with w_inh_ = 1/360, *β* = 1, and *τ* = 50 ms. The external stimulus to each neuron, I_stim,i_, is a Gaussian function, 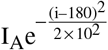. The local excitatory weights are given by a power law function 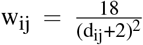 except for Figures 4F,G and 6. T_u_ = 9 and T_d_ = 2 as in the autapse model, except for Figure The stimulus amplitude values simulated are I_A_ = 8, 16, 24, or 32 in Figure 4C and increase with a stepsize of 0.1 from 0 to 60 in Figure 4D.

The time course of each simulation was as follows: With all dendrites originally in the down state, I_stim,i_ was applied for 1000 ms (the ‘encoding period’). The stimulus was then turned off, and the subsequent ‘memory period’ activity was simulated for 1000 ms.

The approximate multistable band of network feedback shown in Figure 4B corresponds to the neuron i = 180 at the center of the bump of activity that determines the height of the activity bump. The B_u_ and B_d_ curves that define the edges of the band were obtained by simulation as follows:

- Tracing the up-threshold curve B_u_. During the encoding period, we plotted the total recurrent feedback 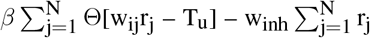 in Eqs. (13) and (14) against the firing rate r_i_, with time as the parametric variable. This numerically generated the up-threshold curve B_u_. The stimulus amplitude used to generate this plot was I_A_ = 80.
- Tracing the down-threshold curve B_d_. The firing rate was first allowed to reach its stable, steady-state value of memory activity. To trace B_d_, we then applied a uniform inhibition, I_stim,i_ = –10 for 1000 ms, until the network activity returned to zero.

As in the autapse case, the amplitude of the decay term, r_i_ in Eq. (13), is represented by a line of unity slope.

In Figure 5A, we used stimulus amplitudes of I_A_ = 20, 40, and 80. In Figure 5B, we applied two successive stimuli centered at the same location. First, a smaller stimulus I_A_ = 20 was applied, leading the subsequent memory activity to stabilize at the location of the red dot. Then a larger stimulus with I_A_ = 80 was applied, followed by a second memory period.

**Figure 5:**
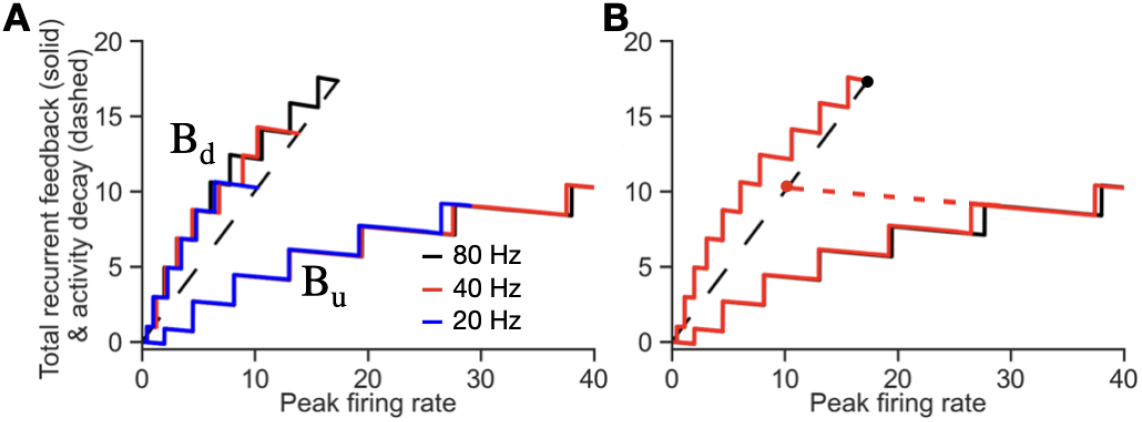
Similar multistable bands under different stimulus amplitudes and presentation patterns. To demonstrate the fidelity of the approximate analytic characterization of the ring model dynamics as being determined by a single multistable band independent of the amplitude of the external stimulus, here we show how little the simulated bands vary under a broad range of different external stimulus conditions. **A**, Varying the stimulus amplitude over a broad set of values. **B**, Applying two successive stimuli, with no offset between them. First, a smaller stimulus amplitude of 20 is applied, resulting in a smaller memory amplitude (red dot). Then, a larger stimulus amplitude of 80 is applied, resulting in a larger memory amplitude (black dot). The black multistable band (hidden on the left side of the band by the overlying red band) is for a stimulus amplitude of 80 with no preceding stimulus, i.e. when starting from zero firing activity.

For the analytical results in Figure 4, the external stimulus was a Gaussian with profile 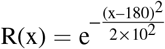, which has an integrated area 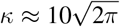.

In Figure 4F,G, simulations were based upon the weight function analytically calculated to produce the linear memory amplitude versus stimulus amplitude relationship M_A_ = c(I_A_–I_A,th_), with c = 0.5 and I_A,th_ = 1.2 (see Mathematical Appendix, Eq. (51)). In Figure 4G, I_A_ takes integer values from 0 to 35.

#### Simulations of networks with noise

For all simulations with noise, spatially and temporally independent Gaussian noise of mean 0 and standard deviation *σ* was applied to the soma of each neuron at each time step of duration dt = 1 ms throughout the entire session.

For Figure 6, the simulated network is based on the one used in Figure 4F. In Figure 6B, *σ* = 3, 6, or 9, and the external stimulus amplitude I_A_ = 10, 15, or 20. 10 trials were conducted for each condition. For each trial, the memory amplitude was estimated by averaging the firing rates of the 10 highest firing neurons for a memory period of 10,000 ms.

**Figure 6:**
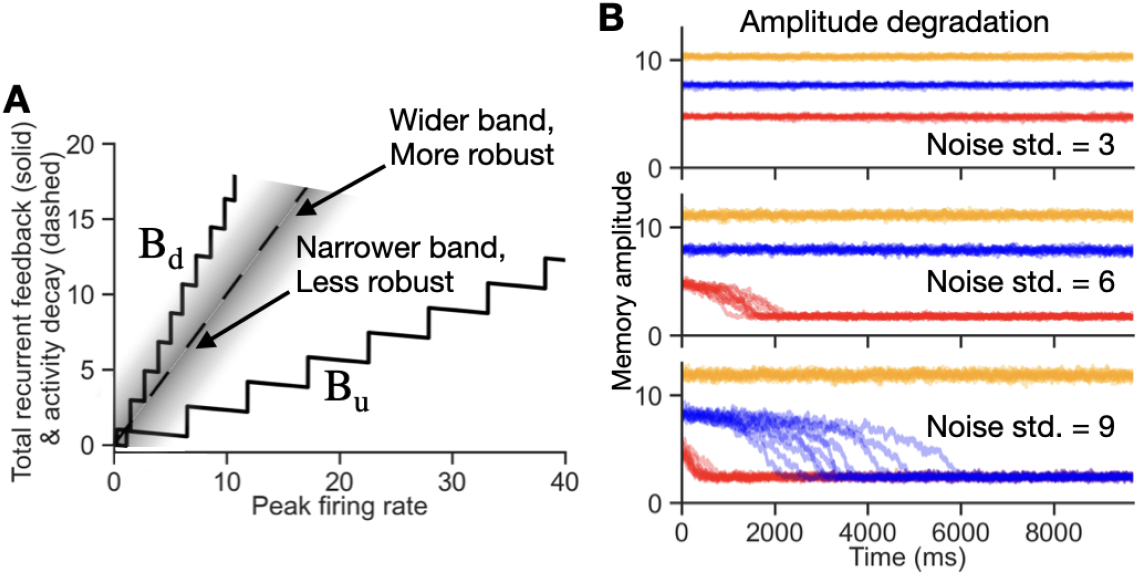
Robustness of memory amplitude to noise and amplitude drift. **A**, Schematic of the effect of noise on memory amplitude. Noise acts as a time-varying external stimulus to each neuron, whose effect can approximately be envisioned as fluctuating the dashed line, along which stable points lie in the no-noise case, horizontally left or right (gray cloud). A memory amplitude is stable if noise does not drive stable memory states across an edge of the band, i.e. if it does not flip dendrites on or off. The robustness to noise depends on the noise level and geometry of the multistable band, with wider portions of the band more robust to noise than narrower portions. **B**, Examples of amplitude drift. Demonstration of the concept sketched in **A** for simulations of the network of Figure 4F with 3 different noise amplitudes (top to bottom panels) for 3 different stimulus strengths (orange, blue, red traces: stronger to weaker stimuli, respectively). Ten simulated trials are shown for each condition, with memory amplitude measured by the average firing rate of the 10 most active neurons in the bump of activity. For small noise (top), all three memories remain stable for each shown stimulus amplitude. For moderate noise (middle), the memory responses with a lower stimulus amplitude (red) go to baseline activity. For larger noise (bottom), memory of the lowest amplitude (red) degrades quickly and that of the middle amplitude (blue) degrades more slowly. The nonzero baseline memory amplitudes following loss of the memory reflect positive fluctuations in firing rate driven by the noise, rather than any remaining storage of memory amplitude.

**Figure 7:**
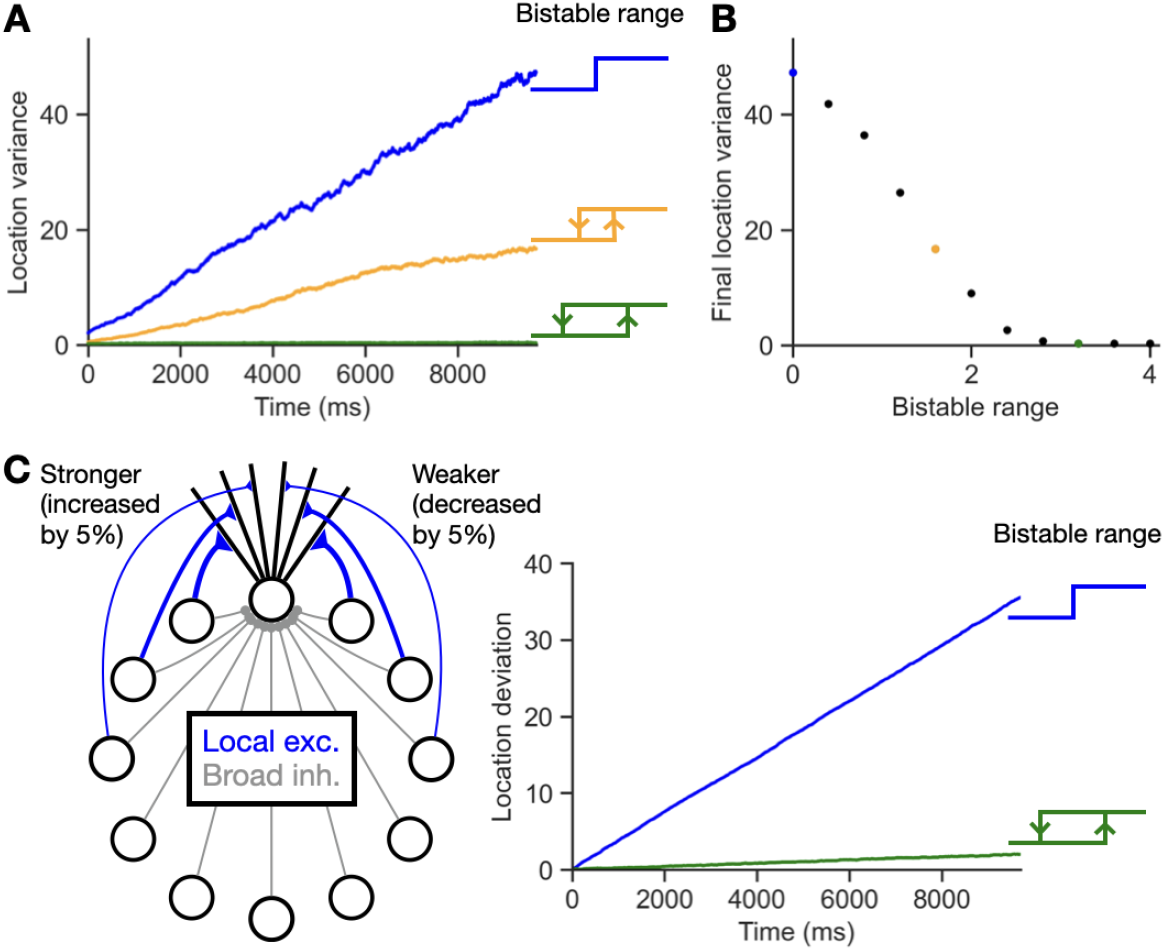
Robustness of memory location to noise with dendritic bistability. **A**, Dendritic bistability reduces the diffusion of the location of the bump of memory activity. Plots show the variance of the memory location over time for 400 trials. If each dendrite has a step function output, T_u_ = T_d_, the diffusion is severe with a large location variance over time (blue). If the dendritic bistable range, T_u_ – T_d_, is larger, the diffusion is reduced (orange or green). For all figure panels, the parameters defining the band of bistability were identical to those shown in Figure 6A except for the values of T_u_ and T_d_ (see Materials and Methods). **B**, Reduction of location diffusion. The variance of the location at the last time step of **A** is recorded for the shown bistable ranges. As the bistable range increases, the location diffusion decreases and eventually becomes negligible when the bistable range is wide enough that noise very rarely affects the dendritic states. Colored points correspond to colored traces in **A. C**, Robustness to asymmetry of ring model recurrent connectivity. Left, the ring model connectivity was perturbed by increasing clockwise directed connections by 5% and decreasing counter-clockwise directed connections by 5%. Right, breaking of the symmetry of the connectivity caused strong drift in the bump location in a model with no dendritic bistability (blue) but very little drift in the model with bistable dendrites (green).

In Figure 7A,B, we simulated diffusion of the bump location under noise for different bistable ranges, starting from the zero-width, step function case with T_u_ = T_d_ and increasing the bistable range T_u_ – T_d_. Increasing T_u_ changes not only the bistable range, but also the strength of synaptic input to each dendrite that is required during the encoding period to achieve a given amplitude of memory activity. Therefore, to approximately keep the memory activity the same for simulations with different bistable ranges, we changed the simulation protocol so as not to start each simulation from zero activity. Instead, we first initialized the network by simulating a network with T_u_ = T_d_ = 3, I_A_ = 15, and no noise for encoding and memory periods of duration 1000 ms. The network activity state at the end of this simulation then served as the initial condition for the shown simulations. From this initial condition, we then tested the effect of applying noise with standard deviation *σ* =10 for a period of 10,000 ms for different values of T_u_ and T_d_. In Figure 7A, the blue trace corresponds to T_u_ = T_d_ = 3, the orange trace has T_u_ = 3.8 and T_d_ = 2.2, and the green trace has T_u_ = 4.6 and T_d_ = 1.4. For each trace, we recorded the variances of the memory locations, estimated by the center of mass of neural activity, and averaged over 400 trials. In Figure 7B, the final location variance is plotted against the bistable range, with T_u_ = 3 + 0.2s and T_d_ = 3 – 0.2s for integers s from 0 to 10. In Figure 7C, we perturbed the connectivity to add a small asymmetric component. This was done by increasing the weights of clockwise-directed connections by 5% and decreasing the weights of counter-clockwise-directed connections by 5%. All other parameters were identical to those used for the blue and green traces in Figure 7A,B.

For Figures 6 and 7, the shown plots begin at 300 ms after the offset of the stimulus to allow the transient remaining effects of the stimulus to decay away.

### Implementations of numerical simulations

For all simulations, we integrated the differential equations using the forward Euler’s method with a time step of 0.01 ms for the spiking model and 1 ms for the rate-based ring model, except for Figures 4B,F, 5, and 6A, where 0.01 ms was used to better approximate the simulated multistable band.

### Code accessibility

Code was written in Python version 3.9.16 and is available in Github [44].

## Results

### Storage of the graded amplitude of a stimulus in an autapse circuit with bistable dendrites

#### Spiking autapse circuit model with multiple bistable dendrites

We constructed a spiking autapse circuit model to demonstrate how a single biophysically based neuron with multiple conductance-based dendritic compartments can robustly sustain graded amplitude activity. The model neuron consists of an integrate-and-fire somatic compartment and 10 conductance-based dendritic compartments (Fig. 2A, Materials and Methods). Spiking output from the soma projects back onto each dendritic compartment through NMDA and AMPA receptor-mediated input, providing recurrent self-excitation. Due to the voltage-dependent activation kinetics of the NMDA receptor and the inward rectifier potassium (Kir) channel, each dendritic compartment exhibits a bistable voltage response to synaptic input, similar to that described previously [43] (Fig. 2B). Briefly, in the absence of synaptic input, the hyperpolarization-activated Kir conductances maintain the dendritic voltage at a strongly hyperpolarized value. As the presynaptic firing rate of the neuron increases from zero, the fast excitatory postsynaptic potentials mediated by the AMPA receptors increasingly activate the slower, voltage-dependent NMDA conductances. Additionally, postsynaptic depolarization reduces the hyperpolarizing effect of the Kir conductance. When the input firing rate crosses an approximate threshold, there is a nonlinear increase in voltage up to a plateau level that is caused by the self-sustaining positive feedback loop between voltage activating the NMDA channels and the NMDA channels increasing the voltage (Fig. 2C, middle). This elevated voltage can then be sustained even for much lower levels of synaptic input. Different dendritic compartments have different synaptic input thresholds for flipping to this elevated, “up” voltage state, with the threshold level determined by the maximal conductance of the NMDA channels (Fig. 2B, dendrites ordered by level of their maximal conductance). The “down” threshold for flipping down the voltage is predominantly determined by the interspike interval at which NMDA-mediated synaptic inputs no longer summate sufficiently to maintain an elevated voltage, and thus is relatively insensitive to the level of NMDA conductance.

Due to the successive recruitment of dendritic compartments with different thresholds, the resulting circuit can sustain many different levels of persistent neural activity that can be used to store a graded representation of the amplitude of a stimulus. When a stimulus is presented (Fig. 2C, top; each pulse represents a separate stimulus), this leads to a transient increase in the firing rate of the neuron. This transient increase in firing then leads to the activation of those dendrites whose activation thresholds are crossed (Fig. 2B). When the stimulus turns off, the activated dendrites can maintain the firing rate of the neuron at a level determined by the number of dendritic compartments activated during the transient stimulus (Fig. 2C). In this manner, the circuit maintains a graded memory of the amplitude of the stimulus, with the number of different amplitude values that can be maintained equal to the number of dendritic compartments. Further, the maintenance of the memory activity is robust due to the bistability of the dendritic response, for which small changes in firing rate cause minimal changes in dendritic voltage (Fig. 2B). In the following section, we use a simplified, firing rate based model to rigorously characterize the core computational mechanism by which this robust storage of graded stimulus values occurs.

#### Firing rate based autapse circuit model with multiple bistable dendrites

We constructed a firing rate based, multi-dendrite autapse circuit model that captures the essence of the spiking model while permitting rigorous, analytic understanding of the memory storing mechanism. The model consists of a neuron receiving input from two sources: an external stimulus I_stim_ whose amplitude is to be stored in the autapse circuit, and N_d_ bistable dendritic compartments that receive recurrent synaptic input driven by the firing of the neuron. The dynamics of the firing rate r obeys:

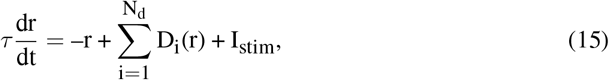

where the first term represents the intrinsic decay of the neuronal firing rate, with time constant *τ*, and the bistable response of each dendritic compartment is given by

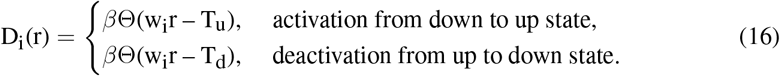

Here, w_i_ gives the strength of the synaptic connection onto the i^th^ dendrite, and *β* gives the contribution of each activated dendrite to the neuron’s firing rate. T_u_ and T_d_ give the threshold values of input to the dendrite, w_i_r, required to flip the dendrite to its up or down state, respectively. Equivalently, the dendritic input-output relation can be characterized by the threshold values of the firing rate required to activate or deactivate the dendrite, 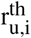 and 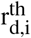 (Fig. 3A, left). Summing together the individual bistable input-output relations of each of the dendrites gives a multistable, staircase-like relation for the total dendritic feedback to the neuron, with the corners of the stairsteps defined by the set of 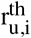 and 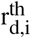 values (Fig. 3A, right; Fig. 3B, gray boxes).

For the neuron to persistently maintain firing activity in the absence of a stimulus (i.e., have dr/dt = 0 when I_stim_ = 0), the recurrent feedback provided through the dendrites must offset the intrinsic leakiness of the neuron. Mathematically, from Eq. (15), this occurs when the total dendritic feedback equals the intrinsic neuronal decay term, 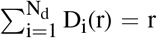. Graphically, this can be depicted by plotting each of these terms as a function of the firing rate of the neuron (Fig. 3B). The total dendritic feedback term 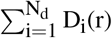 is represented by a multistable band extending between left and right edges B_d_(r) and B_u_(r), respectively (Fig. 3B, rectangles). The intrinsic decay of the firing rate (corresponding to the term –r in Eq. (15)) is given by a line of unity slope (Fig. 3B, blue). The points of intersection of the multistable band and this line (Fig. 3B, blue dots) give the possible stable values of persistent firing activity that can be used to store different stimulus amplitudes. As can be seen, the occurrence of many possible stable values of firing activity does not require fine tuning – due to the width of the bistable band, weights could be changed over a wide range, warping the band, without losing the points of intersection [13, 28, 29]. This contrasts with classic graded amplitude circuits lacking bistable dendrites in which the feedback *band* collapses into a feedback *line* so that fine tuning is required for the circuit to be able to maintain many different persistent firing rates [1].

For the circuit to encode a stimulus into memory, dendrites activated by the external stimulus must remain activated during the subsequent memory period. Graphically, the ability of an external stimulus to activate dendrites can be depicted by shifting the unity-slope line that represents the decay term –r in Eq. (15) to the right by an amount I_stim_ (Fig. 3C, rightward pointing red arrow). Dendrites are activated up to the point where the multistable band first intersects the shifted unity-slope line, with the rate of increase of firing activity proportional to the difference between the edge of the multistable band and the shifted line (Fig. 3C, dr/dt). Subsequently, when the external stimulus is turned off, the system relaxes to a nonzero value (Fig. 3C, blue dot) that reflects the amplitude of the previous stimulus.

The shape of the multistable band is determined by the distribution of synaptic weights w_i_ in the circuit. For a neuron with many dendrites, this weight function can be approximated by a continuous function w(x), where the continuous value x replaces the discrete dendrite index i (Materials and Methods). Different weight functions give rise to different shapes of the multistable band (Fig. 3C-E). These different band shapes result in different relationships between the external stimulus amplitude and the amplitude of memory-storing activity (Fig. 3F). For weight functions with a power law decay of the form w(x) = a/(x + b), as in Figure 3B,C, the band edges are linear, giving rise to a linear relationship between the amplitude of memory activity and the stimulus amplitude (Fig. 3F, blue; Mathematical Appendix). For the weight function used in Figure 3D, the relationship between the stimulus amplitude and memory activity is continuous but with a nonlinear, concave shape (Fig. 3F, gray). As a result, the circuit devotes a relatively higher resolution in memory (larger fraction of its firing rate range) to smaller stimulus values, which could be useful if a task demands more sensitivity to lower amplitude stimuli. In Figure 3E, the stronger nonlinearity in the band shape leads to a discontinuous jump in the memory amplitude at a threshold level of the input stimulus, followed by a very robust, largely saturated response (Fig. 3F, purple). A feature of this connectivity structure is that it amplifies small stimulus values into large firing rates during the memory period. General mathematical formulas for the shape of the multistable band and the stimulus amplitude-memory amplitude relation, for a given weight function, are derived in the Materials and Methods and Mathematical Appendix.

### Working memory in the rate-based ring model with bistable dendrites

Building on the insights gained from the autapse model, we constructed a rate-based ring model consisting of neurons with multiple bistable dendrites. As in classic ring models, neurons in the model are connected through local excitation and broader inhibition (Fig. 4A, left). Each of the N neurons in the network receives recurrent excitatory inputs through bistable dendrites (Fig. 4A, right) and recurrent inhibitory and feedforward stimulus inputs through direct connections onto the soma. The resulting equation for the firing rate r_i_ of the i^th^ neuron in the model is:

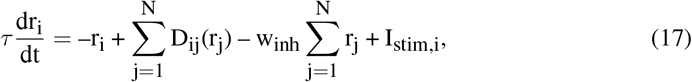

where, as in the autapse model, the first and last terms on the right-hand side represent the intrinsic decay of the neuron and the external stimulus input into this neuron, respectively. The third term corresponds to recurrent inhibitory input not present in the autapse model. This was modeled as being all-to-all with strength w_inh_, but the same qualitative behavior can be obtained with non-uniform lateral inhibition. The second term represents the recurrent excitatory inputs and was governed by dendritic input-output relations of the form:

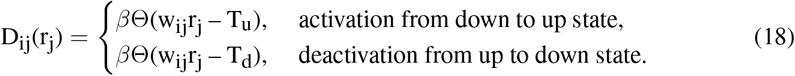

Here, w_ij_ gives the strength of the input arriving onto neuron i from neuron j, which we assume projects to the j^th^ dendrite of neuron i. This assumption that each presynaptic excitatory input arrives on a devoted dendrite was made primarily for analytic tractability, but could be realized biophysically if each of the dendritic compartments in the model corresponds to a dendritic spine. Although the model is written formally as if each neuron has N dendritic compartments, compartments with synaptic strengths w_ij_ too weak to activate the corresponding dendrite could be removed with no change in model output. For the model instantiations considered below, the resulting effective number of dendritic compartments was between 15 and 20.

Intuitively, this model can be seen as a spatially extended version of the autapse circuit model. The autapse circuit directly provides feedback onto its multiple bistable dendrites so as to be able to maintain stable firing at multiple levels. Analogously, in the ring model, a group of active neurons together provide feedback onto themselves to maintain a pattern of neuronal firing that can achieve multiple stable levels. Further, as in traditional ring models, the symmetry of the model’s connectivity implies that it can stably maintain such a multistable bump of activity at any given location. This intuition, and the close relation of this network’s behavior to that of the autapse circuit model, is formalized more quantitatively below and in the Mathematical Appendix.

For activity to be maintained in memory following the offset of a transiently presented stimulus, each neuron must achieve a stable, nonzero firing rate (dr_i_/dt = 0) in the absence of external stimulus input (I_stim,i_ = 0). Similar to the autapse model, this requires that the total recurrent feedback onto each neuron (second and third terms of Eq. (17)) balances the intrinsic decay of the neuron’s firing rate (first term of Eq. (17)). This can be depicted graphically for each neuron by plotting the total recurrent feedback as a function of the neuron’s firing rate (Materials and Methods). As in the autapse model, the feedback forms a multistable band, with left and right band edges defined as B_d_ and B_u_, respectively (Fig. 4B). The negative slope of each step of the total recurrent feedback band is due to the recurrent inhibitory input that was not present in the autapse model. For a given neuronal firing rate and level of steady state recurrent feedback, persistent activity can be maintained by this neuron at the point at which the recurrent feedback band intersects the unity-slope line corresponding to the intrinsic decay of firing (Fig. 4B, dashed line, corresponding to the term –r in Eq. (17)).

A network simulated with this architecture can successfully maintain memory activity that reflects the amplitude of an externally presented stimulus. During the encoding period, when the stimulus is present, neural activity reflects both the direct influence of the external stimulus and the recurrent feedback mediated by the bistable dendrites and recurrent inhibition (Fig. 4C, left, shown for a stimulus that entered the network through a Gaussian shaped input profile). Following the offset of the stimulus, the recurrent interactions between the neurons maintain activity at a firing rate that reflects the amplitude of the external stimulus (Fig. 4C, right). The relationship between the amplitude of the stimulus and the amplitude of the memory activity encoding this stimulus is shown in Figure 4D (blue line).

The graphical construction described above (Fig. 4B) only characterizes the dynamics of a single neuron at a time, rather than the collective behavior of the network as a whole. Because the shape of the band of a given neuron depends on the activity of the other neurons in the network, this single-neuron band is generally insufficient to describe the whole-network behavior in a manner comparable to the analysis of the autapse circuit. However, with some simplifying assumptions, we found that we could define a single multistable band that well-approximated the behavior of the entire network and enabled us to predict the memory amplitude vs. stimulus amplitude relation (such as that in Figure 4D) for a given set of network parameters. As described more fully in the Mathematical Appendix, if we assume that the responses of the network during the encoding period and the memory period are approximately the same shape as that of the external stimulus profile, then the equations for the amplitude of the bump of activity reduce down to equations mathematically analogous to those for the amplitude of activity in the autapse model, modified by a factor accounting for the presence of recurrent inhibition. The resulting analytically calculated band for the whole network has a shape (Fig. 4B, gray lines) that closely approximates the simulated multistable band for the neuron at the peak of the bump of activity (Fig. 4B, black lines), up to deviations that occur at the very highest sustainable firing rates (around approximately 15 Hz in Figure 4B), due to the activity bump notably changing shape for very large stimuli. Away from this saturating limit, the analytically calculated memory amplitude vs. stimulus amplitude relation for the network (Fig. 4D, gray line) closely approximates that of the network simulation (Fig. 4D, blue line). Thus, the same fundamental mechanisms that precisely describe the amplitude-storing dynamics of the autapse circuit can also, to a very good approximation, be used to describe the amplitude-encoding dynamics of the ring model.

The analytic calculations enabled us to determine how the shape of the network’s multistable band and the shape of the memory amplitude vs. stimulus amplitude relationship depend on the synaptic connectivity profile w(x) and the shape of the external stimulus profile R(x), where x is the distance between neurons along the ring (Mathematical Appendix). For example, Figure 4F compares the simulated (black) and analytically calculated (gray) bands for a weight profile derived to produce a linear-shaped band, and Figure 4G shows the resulting memory amplitude vs. stimulus amplitude relationship (gray: analytically calculated; blue: simulations). We note that the analytic model predicts that there is little dependence of the multistable band on the amplitude of the external stimulus (with small deviations resulting from the shape of the bump notably changing for large external stimuli). The simulations verified this prediction, showing that the shape of the band for a given network is quite consistent across a wide range of stimulus amplitudes (Fig. 5A). Further, if a second stimulus is applied while a first stimulus is still being stored, the resulting band for the second stimulus is minimally affected by the first stimulus (Fig. 5B, black is for a single large stimulus and red is for a small stimulus as in Figure 5A, followed by the large stimulus).

### Robustness of the rate-based ring model with bistable dendrites

Traditional continuous attractor models for the storage of the amplitude [1, 2] or location [3, 4] of a stimulus both require fine tuning of their connectivity to maintain analog representations and exhibit a diffusion of activity in the presence of noise that degrades the fidelity of the memory over time [45, 46, 47, 48]. By contrast, model networks with bistable components can have greater robustness to perturbations in their storage of amplitude [13,28,29] or location [30] information. Here, we examine the robustness of the ring model with multiple bistable dendrites to the simultaneous storage of both memory amplitude and location.

We first considered maintenance of the amplitude of a memory state in the presence of noise (Fig. 6). Mathematically, noise appears as a fluctuating external stimulus term in Eq. (17) for each neuron of the network. Thus, similar to the stimulus-driven external input (cf. Fig. 3C), the effect of such noise can be thought of as horizontally jiggling back and forth the dashed line along which stable memory states lie (Fig. 6A). Graphically, this suggests that the width of the multistable band determines the level of robustness to noise: if the standard deviation of the noise input (Fig. 6A, gray shadowing around the dashed line) is much less than the distance to either edge of the band, the memory amplitude is expected to remain stable, whereas if the standard deviation exceeds the distance to either edge of the band, then dendrites will flip on or off, and the network activity is expected to change. Specifically, if the left edge of the band is closer to the no-noise memory state locations (along the dashed line) than the right edge of the band, as in Figure 6A, then noise will more often turn off dendrites than turning them on, and the overall activity will be expected to decline over time. To test this idea, we injected a random white noise input to each neuron in the network and then monitored how well the network could maintain three different external stimulus amplitudes (Fig. 6A, colored lines). For moderate levels of noise, the resulting amplitudes of persistent neural activity are held nearly perfectly constant in memory, exhibiting no diffusion (Fig. 6B, top). When the noise is increased, the lowest amplitude memories degrade (Fig. 6B, middle and bottom) whereas the largest amplitude stimulus could still be robustly stored in memory.

We next considered the maintenance of the location of a stimulus in memory in the presence of noise or asymmetry in connectivity (Fig. 7). For a network without bistable dendrites (Fig. 7A, blue), the variance of the location of the bump of activity in the network in the presence of neuronal noise increased rapidly and approximately linearly over time, consistent with the well-documented diffusion of the memory location over time in classic bump attractor networks [45, 46]. By contrast, for networks with bistable dendrites, the diffusive drift was decreased or nearly completely eliminated (Fig. 7A, orange and green traces), with the amount of diffusive drift decreasing monotonically with the width of the bistable dendritic input-output relation (Fig. 7B). Further, networks with bistable dendrites (Fig. 7C, green trace) could quite stably maintain a bump of activity at a given location in the presence of small asymmetries in network connectivity, whereas the same amount of asymmetry led to rapid drift in the bump location for networks without bistability in their dendrites (Fig. 7C, blue trace). These results reflect that the width of each dendrite’s bistable relation determines its tolerance to noisy input or to perturbations of weights that alter input currents, both for storing the amplitude (Fig. 6) and location (Fig. 7) of a stimulus. Thus, neurons with bistable dendritic compartments can provide a network with a highly noise-tolerant substrate for working memory storage.

## Discussion

Biological systems are capable of remembering both the amplitude and location of stimuli. Previous working memory models have either been capable of storing only amplitude or only location information, or have been non-robust to noise and perturbations of synaptic weights. Here we have demonstrated how amplitude and location may simultaneously and robustly be encoded through the inclusion of bistable dendritic compartments in neurons. For a sufficiently large number of bistable dendrites in each neuron, the model can robustly maintain an approximately continuous amplitude at any given location. We show through simulations and analytic calculations how this robust storage of amplitude information can be understood geometrically as resulting from a multistable band of activity within which activity can be robustly maintained without fine tuning, and how the shape of this band is determined by the form of the synaptic weight distribution.

### Biophysical implementation of the model

The fundamental building block that provides robustness to our model is dendritic bistability. Following previous work [41, 42, 43], we show how such voltage bistability can be realized through the voltage-dependent activation properties of NMDA receptors and Kir conductances. The key feature of these ionic conductances is their voltage-dependent activation, which mediates positive feedback in the cellular voltage dynamics: the depolarization-dependent activation of an inward conductance such as the NMDA channel robustly maintains the elevated voltage state, while the hyperpolarization-dependent activation of an outward conductance such as the Kir channel robustly maintains the low-voltage state. Other ion channels such as voltage-sensitive Ca^2+^ conductances also have this property and have been demonstrated to contribute to cellular or dendritic plateau potentials [32, 40, 49].

NMDA receptors particularly have been demonstrated to be critical to working memory function [50, 51, 52] and have been a major component of working memory models [12]. However, it is still unclear whether this is simply due to their slow kinetics, which makes memory activity more stable [53], or whether it is also related to their ability to induce dendritic bistability. Physiological recordings have shown that NMDA receptors in dendrites can cause prolonged voltage plateaus lasting hundreds of milliseconds [49, 54, 55], but more direct evidence is needed to determine whether dendritic plateau potentials can be sustained for seconds in the presence of persistent synaptic input during working memory tasks. Even if dendritic compartments only sustained prolonged plateau potentials, rather than maintaining truly bistable states, such plateau potentials could nevertheless provide a long intrinsic timescale for working memory that is then extended by network-level feedback interactions [2, 13].

A critical question is how many bistable states can be independently maintained in a single neuron. Classic modeling work based upon morphologically reconstructed neurons [56, 57, 58, 59] suggests that neurons are capable of maintaining several tens of approximately independent voltage compartments, consistent with experiments suggesting highly compartmentalized function [49, 55, 60, 61, 62, 63]. Very many compartments could be maintained independently if the functional voltage subunit was a dendritic spine, as suggested by recent experiments [62, 63], with localization of bistable voltage compartments enhanced by the conditionality of NMDA receptor activation on presynaptic input activity [41].

Alternatively, the multistable single-cell dynamics studied here could be realized in a larger network in which either individual neurons [34, 35, 36, 37, 38, 39, 40] or clusters of neurons [13] provided the fundamental bistable subunit. In this case, the successive recruitment of bistable dendrites in our model would need to be replaced by the successive recruitment of bistable neurons connected in an appropriate manner to functionally produce the stable, graded-amplitude activity demonstrated here. For example, a literal mapping of our model to a network architecture could be generated if, instead of dendrites projecting to a soma, neurons projected to a common neighbor that acted like a hub, with this hub neuron then providing output to other, similarly arranged neuronal groups arranged in a ring-like architecture. Such an architecture would presumably need to be learned through training. Towards this direction of learning connections between bistable units, recent work inspired by gated recurrent units (GRU) from machine learning have suggested how the synaptic connectivity of large networks of bistable units may be trained to solve tasks requiring memory to be stored in neural activity over long-timescales [64].

### Robustness to noise and sensitivity to external stimuli

Our model shows a trade-off between the sensitivity to small stimuli and the robustness to noise. A wide bistable range in dendrites makes memory robust to noise in neural activities. However, this comes at the cost of reduced sensitivity to the encoding of small external stimuli [28], as a small stimulus may not be able to flip on dendrites and thereby encode information into memory.

This trade-off between robustness to noise and sensitivity to small stimuli might not be inevitable. At least in experiments studying the maintenance of head direction coding in Drosophila [65, 66], small head velocity stimuli can change the head direction value that is stored in memory. One possibility is that earlier stages of processing transform small stimuli to sufficiently large amplitudes that they can be encoded in the network [67]. Similarly, if spikes from external stimuli arrive in a correlated or synchronous manner, then each correlated spiking event corresponding to a stimulus may be sufficiently large to be encoded in memory. This would require that noise, by contrast, arrives in an uncorrelated manner, so that it does not disrupt encoding by bistable dendritic units. Alternatively, the noise robustness/small stimulus sensitivity tradeoff might be avoided if the dendritic compartments can flexibly switch between bistable and non-bistable input-output relations. This might be accomplished if, for example, the external stimuli to a network not only provide excitatory inputs to be encoded by the network but also simultaneously mediate disinhibition of dendritic compartments, changing their current-voltage relationships to a non-bistable regime. Such disinhibition enabling the occurrence of local regenerative dendritic events has been observed to occur in the hippocampal CA1 region, with lateral entorhinal cortical inputs providing disinhibition of CA1 pyramidal neuron dendrites [68, 69]. Altogether, these mechanisms suggest how networks may be able to take advantage of the robustness to noise and perturbations provided by bistable processes while also remaining sensitive to small stimuli.

### Encoding of stimulus amplitude, width, or uncertainty in the multistable ring network

Previous ring models have primarily focused on the encoding of location information while neglecting the ability of networks to encode graded information at a given location. Such graded amplitude encoding is the essence of line attractors for parametric working memory. However, line attractors generically require fine tuning to generate the continuum of fixed points along the attractor basin [2, 3, 5, 13]. A small set of previous studies have produced ring networks that successfully maintain amplitude information [23, 24, 25, 26], but like classic line attractors, these networks all involved a fine tuning of synaptic connectivity. Alternatively, a recent study showed that a discrete set of amplitudes could be encoded in a ring network with oscillatory activity [27], but this network had a tradeoff between the degree of required tuning and the discreteness of the activity states, such that storing a larger set of amplitude levels required a larger degree of fine tuning. Building on previous work constructing robust, not fine-tuned line attractor models through the use of multiple bistable units [13, 28, 29], we have shown how the same principles enable the construction of a network that robustly stores both amplitude and location information. In addition, as in previous ring models with cellular bistability [31], our model can store stimuli of different widths, because the width of the sustained bump of activity in our models approximately corresponds to the width of the external stimulus (Fig. 4C and Mathematical Appendix).

The amplitude of a pattern of neural activity can be interpreted as encoding information in two different manners: as the intensity of a stimulus at a given location or as the precision of the location. Here, we have largely framed our discussion of the ring model in the former manner, as encoding the location and intensity of a stimulus. Alternatively, Bayesian models of sensory coding [26,70,71,72,73] show how the amplitude of neural firing can represent the precision of a location, with higher firing rates indicating higher precision. Such a representation of precision enables Bayesian computations to be performed by summations of firing rates [19,20]. Building on this Bayesian framework, recent work has shown how a ring model with slowly decaying amplitude of activity can encode uncertainty about location or orientation [26]. This encoding of uncertainty enabled the network to perform accurate Bayesian inference in tasks such as path integration and evidence accumulation, with suggestions for how this form of coding may apply to the head direction system. Within the Bayesian framework, our dendritic multistability model provides a mechanism by which uncertainty about location or orientation could be robustly maintained in working memory. Thus, dendritic bistability could provide an intrinsic cellular mechanism that enhances the robustness of computations underlying both working memory and Bayesian inference.

### Mathematical Appendix

Here we provide the analytical derivations that underlie the exact or approximate characterizations of the dynamics and steady-state responses of the rate-based model networks. First, we present the calculations that exactly relate the memory-storing properties of the autapse circuit to its weight function. Next, for the ring model, we use a simplifying assumption to derive its approximate multistable band. We then show how, under this simplifying assumption, the mathematical form of this band and the relation between the memory amplitude, stimulus strength, and weight function reduce to a form similar to that of the simpler autapse model.

### Determination of the relationship between the memory amplitude, the stimulus strength, and the weight function in the rate-based autapse model

Here, we derive relationships between the steady-state amplitude of memory activity r_m_, the stimulus strength I_stim_, and the weight function w(x). Specifically, below we derive: (1) the form of the weight function w(x) required to achieve a given stimulus-memory amplitude relationship,

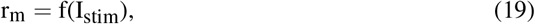

and (2) for a given weight function w(x), the function f().

To proceed, we make two assumptions. First, we assume that the neuronal activity during the encoding period is monotonically non-decreasing, so that dendrites can only remain at their previous value or become activated, and that the activity reaches steady-state by the end of the encoding period. Thus, following the derivation of Eq. (12), the total number of dendrites, z_tot_, that are active at the end of the encoding period is given by

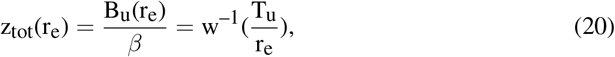

where r_e_ represents the firing rate at the end of the encoding period. Second, we assume that the down threshold T_d_ is low enough that no dendrites flip from their up to down state following the offset of the external stimulus, i.e. that the B_d_ edge of the band is always to the left of the unity-slope line along which fixed points of the dynamics lie (Fig. 3B-E). Thus, the steady-state activity during the memory period differs from that of the encoding period, r_e_, only by removal of the external stimulus I_stim_:

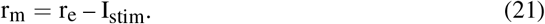

Substituting Eq. (20) into Eq. (21) then gives a relationship for the weight function in terms of the memory period activity r_m_:

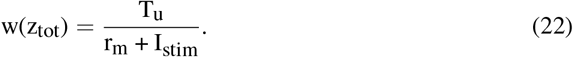

The steady-state memory period activity r_m_ is given by *β* times the number of activated dendrites,

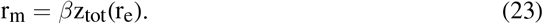

Substituting the above equation and the desired relationship Eq. (19) into Eq. (22) then gives:

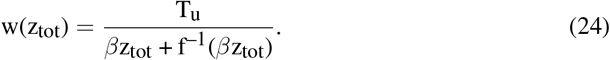

Since this holds for any number x = z_tot_ of dendrites that can be activated by an external stimulus of amplitude I_stim_, we finally have an expression for the weight function required to produce a given relationship f() between the stimulus amplitude and memory amplitude:

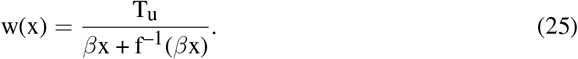

The relations above can be rearranged to determine the relationship f() between the stimulus amplitude and memory amplitude for a given weight function w(x). From Eqs. (22) and (23),

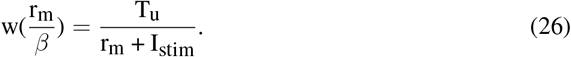

Thus, for a given weight function w(x),

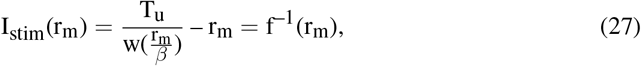

and f() is given by the inverse of this relationship.

#### Weight function for a linear memory-stimulus relationship and linear multistable band

For a linear stimulus amplitude-memory amplitude relationship r_m_ = c(I_stim_ – I_th_), we have that:

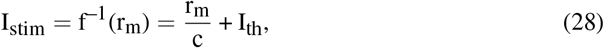

which from Eq. (25) gives:

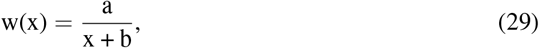

where 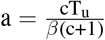 and 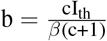. This relationship is shown in Figure 3F for a = 4 and b = 1. For this case, as shown in Figure 3C, the edges of the multistable band are also linear, as may be confirmed by substituting the above form of w(x) into Eq. (20). For example, performing the calculation for the right side of the band gives:

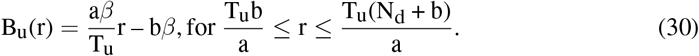

### Derivation of the approximate dynamics of the rate-based model with ring architecture

In this section, we provide analytic calculations of the approximate dynamics of the ring model. Unlike the autapse circuit, it is difficult to provide a simple, exact analytic characterization of the system dynamics described by Eq. (13). This is because, although each individual neuron’s behavior for a given location of the memory-storing bump can be qualitatively conceptualized by a multistable band defined by the sum of its dendritic and recurrent inhibitory inputs, each dendrite in the ring model receives input of a different firing rate r_j_. Therefore, the shape of such a multistable band does not just depend on the local excitatory weights w_ij_, but also on the specific pattern of firing rates r_j_. Thus, because r_j_ depends on the amplitude and shape of I_stim,i_, the exact shape of the multistable band is not uniquely defined. However, as shown in the following sections, we can analytically derive an approximate multistable band (Fig. 4B,F) and relationship between stimulus amplitude and memory amplitude (Fig. 4D,G) that closely parallel the exact results from the autapse circuit and that well-approximate the corresponding simulation results.

#### Calculation of how the shape of the weight function determines the multistable band and the mapping between the external stimulus and memory amplitude in the ring network

Below, we first derive an approximate shape of the multistable band. We then determine approximate relationships between the amplitude of the external stimulus, the memory amplitude (i.e. steady-state firing rate of the neuron at the peak of the activity bump during the memory period), and the weight function. In particular, we show below how the derived relationships tightly parallel and generalize those found for the autapse case, enabling the intuitions derived exactly for the autapse case to be related to the full ring model case.

During the encoding period, we assume for simplicity that neurons start from zero firing rate and only maintain or increase their firing rates when the external stimulus is presented. In this case, the term D_ij_(r_j_) can be replaced in Eq. (13) by a step function, of height *β*, corresponding to the condition for activating a dendrite from its down to its up state:

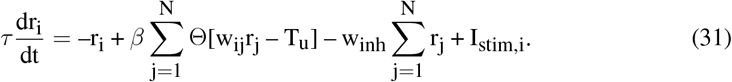

In the following, we work in the continuum limit, with the continuum neuron index x replacing the discrete index i. We denote the steady state firing rate that is approached during the encoding period by r_e_(x), which is given from the above equation by

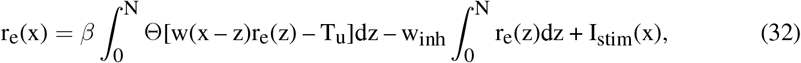

where the excitatory weights w(x – z) are a function of the shortest distance between x and z along the ring. A dendrite is activated if w(x – z)r(z) *>* T_u_. Thus, the first integral gives the total number of activated dendrites. In Figure 4E, we illustrate schematically, for the maximally firing neuron of a bump of activity, how this integral can be graphically solved in response to a localized symmetric stimulus I_stim_(x). The total dendritic input (w(x – z)r(z), black) to postsynaptic neuron x is plotted as a function of the presynaptic neuron index z. Activated dendrites are those (labeled in blue) that exceed the threshold value T_u_ (dashed line).

To derive an approximate multistable band and approximate relationships between the weight profile w(x), external stimulus amplitude, and steady-state memory amplitude, we make two further assumptions. First, we assume T_d_ is low enough that no dendrites flip from their up to down state following the offset of the external stimulus (i.e. during the memory period). Therefore, the steady-state activity during the memory period r_m_(x) is:

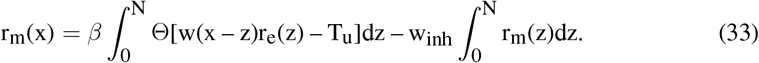

Note that the first integral in Eq. (33) is identical to the first integral in Eq. (32), indicating that the total number of activated dendrites is unchanged. Second, we assume that the stimulus I_stim_(x) and the steady-state firing rates during the encoding period r_e_(x) and during the memory period r_m_(x) approximately have the same symmetric, unimodal (bump-like) shape R(x), so that we can write I_stim_(x) = I_A_R(x), r_e_(x) = E_A_R(x), and r_m_(x) = M_A_R(x). We define R(x) to have a peak value of 1, so that I_A_, E_A_, and M_A_ give the amplitudes of the corresponding stimulus or activity profiles.

#### Deriving the approximate shape of the multistable band

Given these assumptions, we first show that the network activity is governed by a multistable band of recurrent network feedback analogous to that derived for the autapse case (Eq. (12)). We focus our analysis on the neuron at the peak of the bump of activity, which we label without loss of generality as x = 0. From the paragraph above, the firing rates of all other neurons in the network can then be derived by scaling the activity of this neuron by the bump-shape factor R(x). During the encoding period, the total dendritic feedback providing input to neuron x = 0, which we denote as D_tot_(E_A_), is given from Eq. (32) by

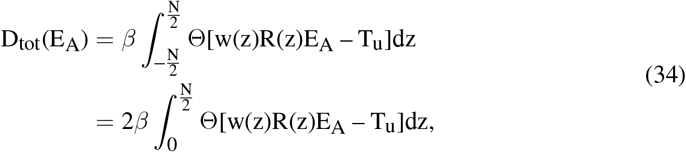

where in the above we have used the symmetry of the weight matrix and the neural activity profile. Performing the integral analogously to the calculation for the autapse case then gives:

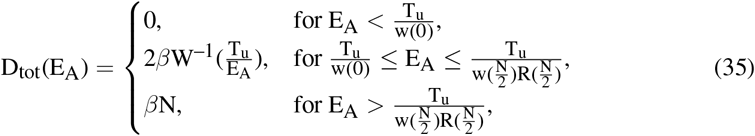

where W^−1^(x) denotes the inverse of the function W(x) = w(x)R(x). Therefore, the total recurrent feedback at steady-state during the encoding period, which specifies the right side of the multistable band as a function of the amplitude of the bump of neural activity, is given from Eq. (32) by

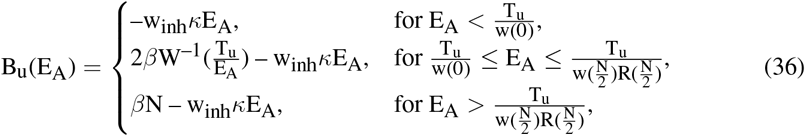

where 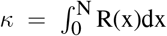 is a constant for a given shape R(z) and depends only on the total integrated area of R(z). Analogous reasoning can be applied to determine the left side of the multistable band, B_d_. In the case of no inhibition, w_inh_ = 0, this result is analogous to that found for the autapse model (Eq. (12)) with W acting analogously to the autapse weight function w and the factor of two being due to the range of x extending symmetrically from negative to positive values for the ring model. This analytical result for the band shape is compared to simulation results in Figure 4B,F.

#### Deriving the relationship between the memory amplitude, the stimulus strength, and the weight function

We next derive the symmetric weight profile w(x) required to generate a given relationship,

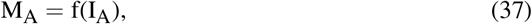

between the external stimulus amplitude and memory amplitude. Our derivation closely parallels that shown above for the autapse model.

First, recall that the memory amplitude M_A_ is defined as the firing rate of the neuron at the peak of the bump of activity during the memory period, and that this peak firing rate is determined by the total number of dendrites, z_tot_, that are active at the end of the encoding period. For the nonzero and not saturated regime of Eq. (35),

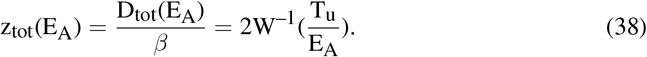

z_tot_ is illustrated as the length of the blue segment in Figure 4E. Defining z_max_ = z_tot_/2 as the index of the last activated dendrite (Fig. 4E, abscissa value of red point), from Eq. (38) we have that:

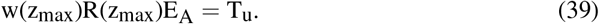

To derive an expression for the memory amplitude M_A_ in terms of the stimulus strength I_A_, first, we show how the steady-state amplitude of the bump at the end of the encoding period, E_A_, is related to I_A_. During the encoding period, and recalling that E_A_ = r_e_(x = 0), Eq. (32) gives:

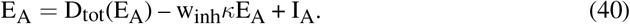

Second, during the memory period, from Eq. (33), the memory amplitude M_A_ is given by the steady-state activity of the neuron at the peak of the activity bump:

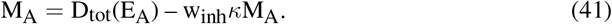

Substituting the expression for D_tot_(E_A_) from Eq. (41) into Eq. (40) and rearranging then gives:

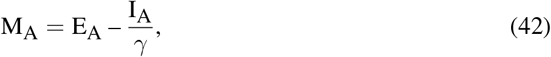

where *γ* = 1 + w_inh_*κ*. This relationship is analogous to Eq. (21) for the autapse model.

Substituting Eq. (42) into Eq. (39) then gives a relationship for the weight function in terms of M_A_:

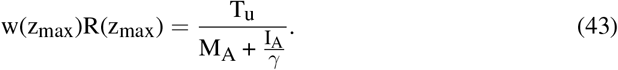

From Eqs. (41) and (38), the memory amplitude M_A_ is related to the number of activated dendrites by

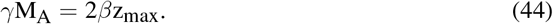

Substituting this equation into Eq. (43) and using that I_A_ = f^−1^(M_A_) gives:

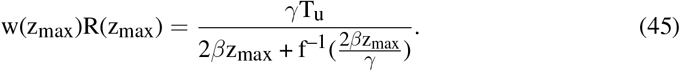

Since the above equation is a requirement on w(x) for any distance x = ±z_max_ from the peak of the bump that could occur with different stimulus amplitudes I_A_, we finally have the general relation:

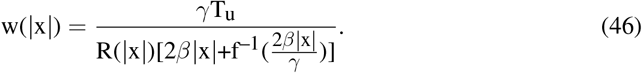

This equation is analogous to Eq. (25) for the autapse model, but modified by a factor *γ* = 1 + w_inh_*κ* that goes to one in the absence of inhibition, a spatial profile term R(|x|), and a factor of two that accounts for the ring model weight profile being defined in terms of negative and positive x values.

The relations above can be rearranged to determine the relationship f() between the stimulus amplitude and memory amplitude for a given weight function w(x). Rearranging Eq. (43) gives:

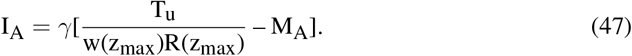

Substituting the expression for z_max_ from Eq. (44) into this relation gives:

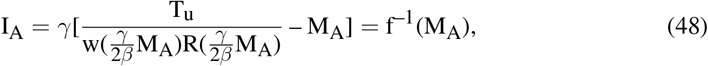

which can be inverted to give f(M_A_). This formula parallels the analogous relation for the autapse case, Eq. (27). In the absence of recurrent inhibition, *γ* = 1, this expression only differs from the autapse case by the spatial profile term R(x).

*Linear memory-stimulus relationship*. For the particular case of a linear relationship,

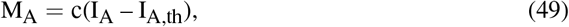

between the amplitude of the external stimulus and the amplitude of the memory period activity bump illustrated in Figure 4G, we have:

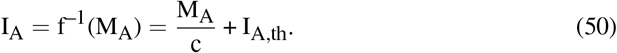

Substituting this relation into Eq. (46) gives:

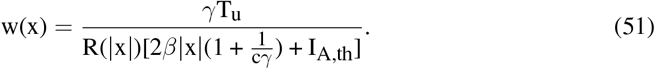

Thus, analogous to the autapse case, the effective weight function W(x) = w(x)R(x) to produce an (approximately) linear input-output function has the form 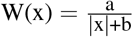. Conversely, for this form of weight function, the slope and threshold of the memory amplitude - stimulus amplitude relationships are given from Eqs. (43), (44), and (50) by

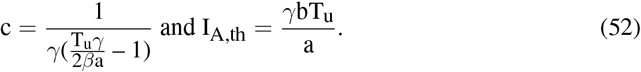

We note that, for a flat stimulus shape, R(x) = constant, this reduces to the formula found for the rate-based autapse case. In addition, note that, although w(x)R(x) decreases monotonically for monotonically decreasing stimulus profiles, w(x) from Eq. (51) may not be monotonically decreasing in |x| or may even blow up if R(x) approaches zero at the edge of the bump. To offset this, and ensure that w(x) is monotonically decreasing as |x| increases, in all simulations, we manually set w(x) to zero once it stopped monotonically decreasing.

## Acknowledgments

This work was supported by the Air Force Office of Scientific Research (AFOSR) award number FA9550-22-1-0532 (JX, DLC) and NIH grants U19 NS132720 and R01 EY027036 (MSG). We thank the Goldman laboratory for helpful discussions and suggestions, and J. Bhasin for critical feedback on the manuscript.

